# CD11B^+^CD36^+^ cells are bone anabolic macrophages that limit age-associated bone loss

**DOI:** 10.1101/2024.09.13.612932

**Authors:** Jinsha Koroth, Ismael Y. Karkache, Elizabeth K. Vu, Kim C. Mansky, Elizabeth W. Bradley

## Abstract

Disruptions in the bone remodeling cycle that occur with increasing age lead to degeneration of the skeleton and increased risk of fragility fractures. Our understanding of how bone remodeling within cortical bone is controlled and altered with age in males and females is limited. Here, we generated bone marrow chimeric mice to understand the impacts of age and sex on bone remodeling. We demonstrate that transplantation of aged male or female bone marrow into young, lethally irradiated male hosts unexpectedly enhances cortical bone mass without impacting cancellous bone. Our single cell RNA-sequencing data show that mice reconstituted with aged bone marrow exhibited subsets of cells marked by CD11B/CD36 expression that demonstrate enhanced production of anabolic cytokines as compared to young counterparts, and that these myeloid subsets exist under conditions of normal physiology in aged mice. Importantly, CD11B^+^CD36^+^ cells do not differentiate into osteoclasts in vitro, and CD36 does not mark TRAP+ cells in vivo. Instead, CD36^+^ cells localize to resorption sites, including within cortical bone defects, suggesting their involvement in cortical bone remodeling and healing. CD11B^+^CD36^+^ cells also express elevated levels of bone anabolic WNT ligands, especially Wnt6. In functional assays, we demonstrate that soluble factors produced by CD11B^+^CD36^+^ cells enhance osteoblast progenitor commitment, mineralization, and activation of WNT signaling in vitro. Moreover, CD11B/CD36 exquisitely mark a subset of anabolic myeloid cells within human bone marrow. In conclusion, our studies identified a novel population of aged macrophages that limit cortical bone loss.

## INTRODUCTION

Bone mass in men and women starts to decline during the third decade of life (1). Women also exhibit a punctuated period of bone loss following menopause proceeded by age-associated declines in bone mass at a rate approaching that of men (2). In both men and women, age constitutes the greatest risk for a fragility fracture (3). Half of all women and one quarter of men are projected to suffer an age-related low bone mass fracture (4). Importantly, approximately 25% of individuals die a year after a hip fracture (5–9); thus, a better understanding of the pathophysiology of age-related declines in bone mass would aid in the development of preventative and interventional strategies.

Cortical bone comprises the majority of skeletal bone mass (80%), and accounts for 70% of bone lost with age (10, 11). Although bone anabolic agents enhance vertebral cortical bone mass, efficacy is lower at the hip (+3.9%) compared to the spine (+12.3%) (12, 13); therefore, additional therapeutics improving cortical bone are desperately needed.

The process of bone remodeling facilitates homeostasis of the adult skeleton. Bone remodeling consists of the activation, resorption, reversal, and formation phases (14). Aging disrupts these phases, resulting in unbalanced bone formation and bone resorption (14). Osteoclasts mediate bone resorption, whereas osteoblasts orchestrate formation. While the coupled cellular activities of osteoclasts and osteoblasts maintain optimal bone health, recent evidence suggests that macrophages within the bone niche help coordinate the bone remodeling process (14). In spite of this, we understand little about how different subsets of macrophages impact the bone remodeling. In this study, we sought to define how age affects macrophage types and examine the impact of those cells on cortical and trabecular bone homeostasis.

## RESULTS

### Transplantation of aged bone marrow into young hosts promotes cortical bone mass

Recent work supports that myeloid cells residing within the bone marrow regulate bone remodeling during conditions of both normal physiology and pathological conditions (15, 16). Lower peak bone mass and acute bone loss following the menopause predisposes women to osteoporosis and greater risk of fracture with age; however, it is unclear if rapid bone loss associated with sex steroid deficiency and age-associated bone loss are mediated by the same mechanisms, or if there are differences in how age-associated bone loss occurs in males and females. To understand the cellular mechanisms driving bone loss prior to reproductive senescence, we performed bone marrow transplantation assays generating chimeric mice. We started with young (e.g., 6-week-old) male recipient mice (Figure 1A, left panel). Young recipient male mice were lethally irradiated to ablate the bone marrow. In parallel, we collected bone marrow from 6-or 40-week-old donor male or female mice (Figure 1A, right panel). Donor bone marrow was depleted of T cells through magnetic-assisted cell sorting (MACS)-mediated removal of CD3+ and CD8+ cells. Young recipient mice were then reconstituted with T-cell depleted donor bone marrow from mice of one of four groups (Figure 1A, right panel) (17–19). We then aged recipient male mice to six-months (e.g., 18-weeks after BM transplantation). To estimate percent chimerism, we collected bone marrow and liver cells from male recipient mice transplanted with female bone marrow, and assessed expression of X-and Y-linked genes. Within the bone marrow, male recipient mice showed robust expression of X-linked genes *Xist* and *Gata1*, but little to no expression of Y-linked genes *Sry* and *Ddx3y* (Figure 1B), supporting that cells within the bone marrow of ablated recipient males were reconstituted by female cells. In contrast, X-and Y-linked genes were expressed within the male recipient liver. Gross examination showed that male mice receiving bone marrow from either aged males (group 2) or aged females (group 4) exhibited signs of premature aging (e.g., gray coat color) (Figure 1C). Body weight was also reduced by transplantation of aged male bone marrow (e.g., group 2), but no other significant weight changes were observed (Figure 1D). Splenomegaly can point to deficiencies in bone marrow function, so we also assessed the spleen weight to body weight ratio. Changes in body weight-adjusted spleen mass were not observed within any group (Figure 1E).

**Figure 1.**
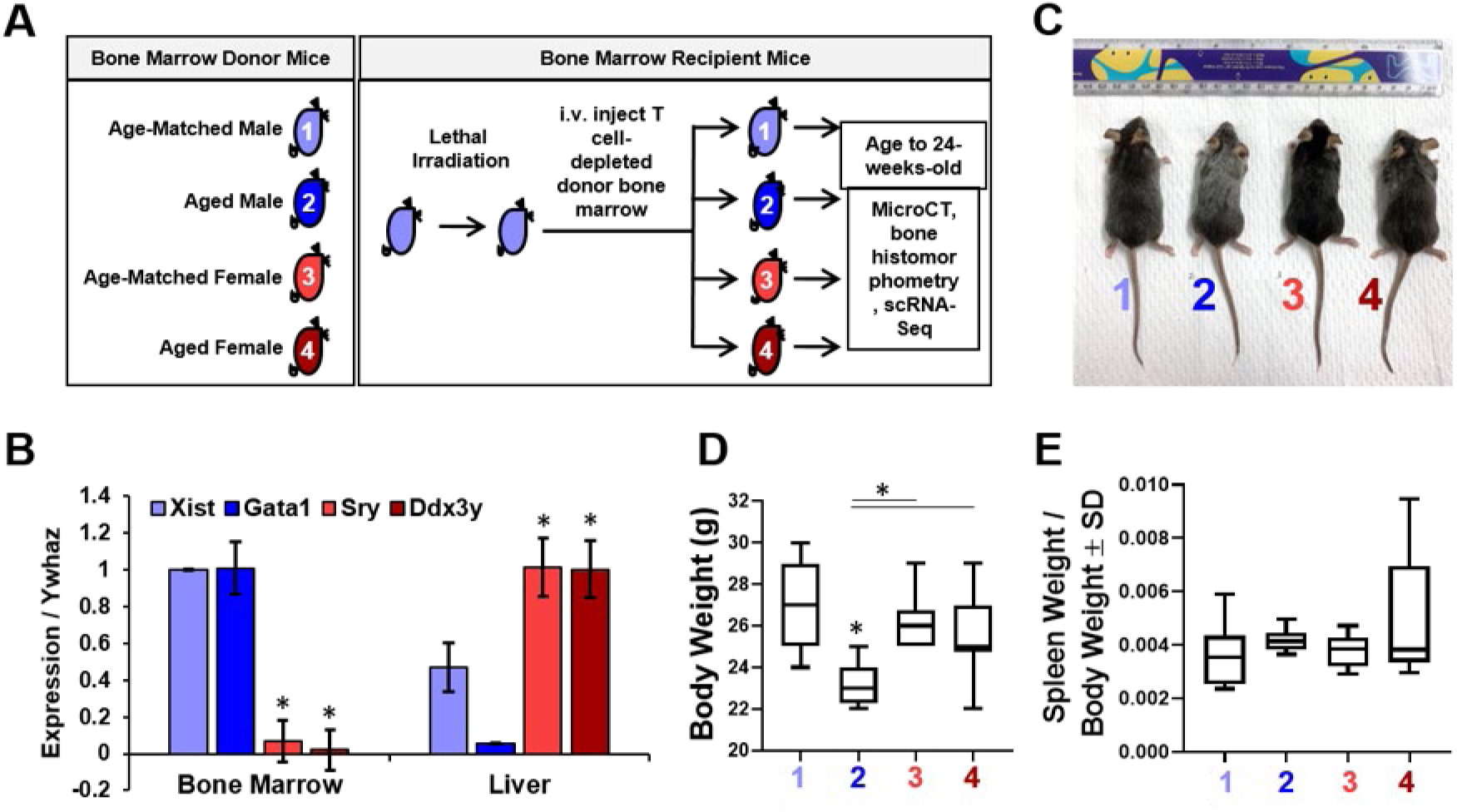
Generation of Bone Marrow Chimeric Mice. (A-E) Young male recipient mice were lethally irradiated and reconstituted with T cell-depleted bone marrow (BM) from 1) young male, 2) aged male, 3) young female, or 4) aged female donor mice. Recipient mice were aged to 24-weeks for subsequent analyses. (A) Experimental overview. (B) RNA was collected from BM and livers of chimeric mice in group 3 and expression of X-and Y-linked genes was determined via qPCR, \**p*<0.05, n=8. (C-E) Chimeric mice in each group were collected six months post-BM transplantation. Mice were (C) photographed. (D) Body and (E) spleen weight/body weight was assessed, \**p*<0.05. Groups: 1: young male BM, n=10; 2: aged male BM, n=8, 3: young female BM, n=8; and 4: aged female BM, n=8

We next assessed bone mass of 24-week-old recipient male mice. We first determined how irradiation affected cortical bone by comparing chimeric mice within each group to 24-week-old non-irradiated controls. We noted that irradiation decreased cortical BV/TV (Figure 2A, D), but did not alter cortical thickness (Figure 2B, D) or trabecular BV/TV (Figure 2D, E) when comparing non-irradiated controls to male mice transplanted with sex-and age-matched bone marrow (e.g., group 1 vs NI). Likewise, enodsteal perimeter was significantly increased when comparing to non-irradiated controls (Figure 2C). In contrast, irradiation increased trabecular spacing but reduced trabecular thickness (Figure 2D, F, G).

**Figure 2.**
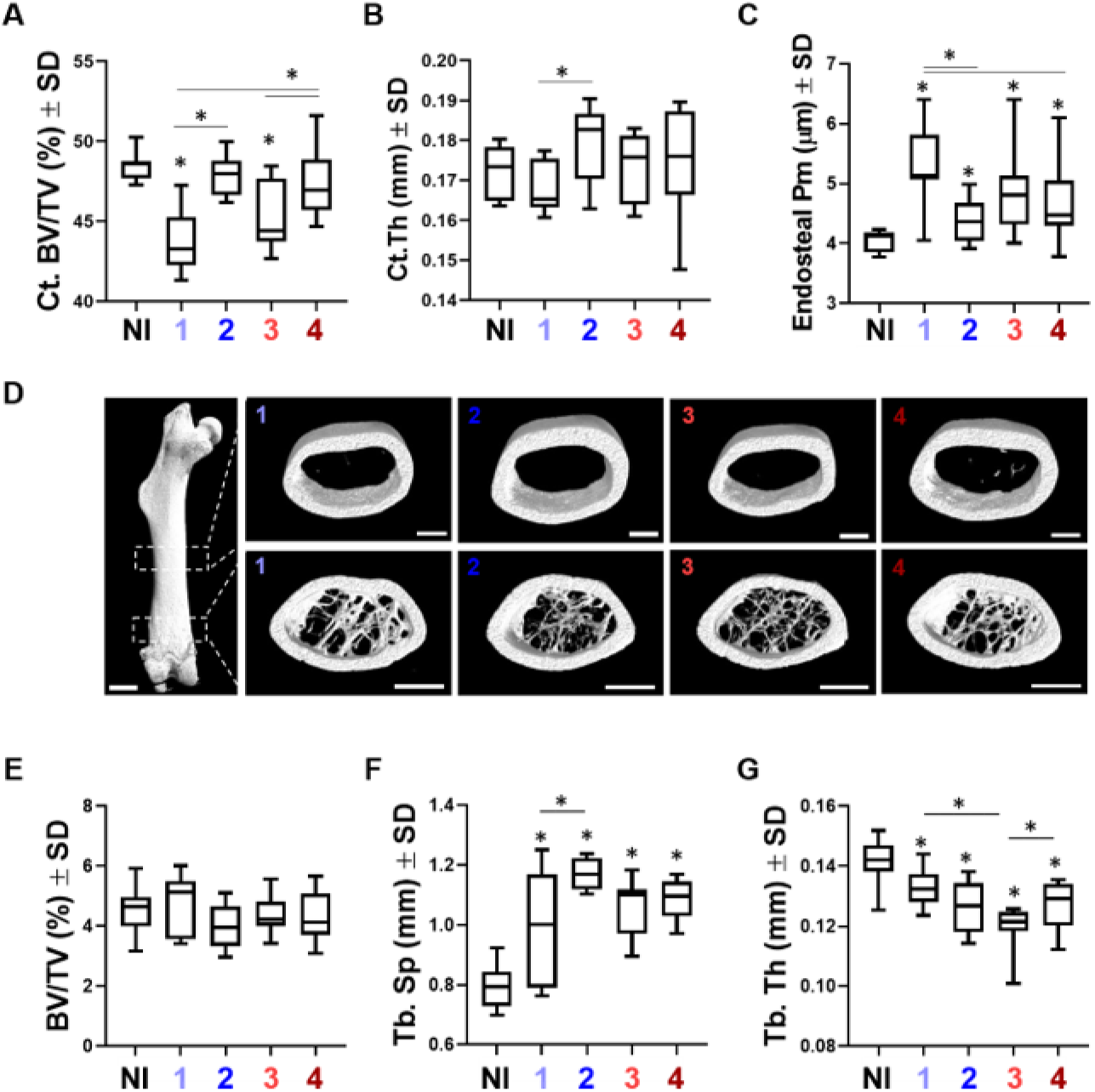
Transplantation of Aged Bone Marrow Enhances Cortical Bone. (A-G) Young male recipient mice were lethally irradiated and reconstituted with T cell-depleted bone marrow (BM) from 1) young male, 2) aged male, 3) young female, or 4) aged female donor mice. Recipient mice were aged to 24-weeks for subsequent analyses. Non-irradiated 24-week-old males were used for comparisons. Micro-CT of the femoral mid-shaft was performed to determine (A) cortical BV/TV, (B) cortical thickness, (C) endosteal perimeter (Pm) and (D) reconstructions were generated. The distal femoral region was also assessed to determine trabecular (E) BV/TV, (F) trabecular spacing, and (G) trabecular thickness. #*p*< 0.05 NI vs group 1, \**p*<0.05 vs. group 1. Groups: NI: non-irradiated control, n=10; 1: young male BM, n=10; 2: aged male BM, n=8, 3: young female BM, n=8; and 4: aged female BM, n=8

We next assessed changes between groups 1 and 2 to determine the effect that aged, but sex-matched bone marrow had on cortical and trabecular bone. Micro-CT analyses of the resulting chimeric mice in groups 1 and 2 revealed a surprising increase cortical bone mass (+10%) and cortical thickness (+7%) when male recipients were reconstituted with sex-matched, but aged bone marrow (Figure 2A, B, D). In contrast, no changes in trabecular BV/TV were observed between these groups (Figure 2D, E), but we did note increased trabecular spacing coupled with diminished trabecular thickness (Figure 2D, F, G). These data suggested that transplantation of aged bone marrow enhanced cortical but not trabecular bone mass.

We next examined the effects transplantation of bone marrow derived from female mice on bone mass. When comparing young male mice reconstituted with bone marrow from age-matched males or females (groups 1 and 3), we did not see a change in cortical BV/TV or cortical thickness (Figure 2A, B, D). Likewise, trabecular BV/TV, trabecular number, and trabecular spacing were unchanged (Figure 2D-F), but we did observe a slight reduction in trabecular thickness when comparing these groups (Figure 2G).

We next compared young male mice transplanted with either sex-and age-matched bone marrow to those transplanted with bone marrow collected from 40-week-old females (groups 1 and 4). We noted increased cortical BV/TV, but no significant changes in cortical thickness (Figure 2A, B, D). No changes in trabecular bone parameters were noted (Figure 2D-G). We then compared the effects of age when transplanting female bone marrow (e.g., groups 3 versus 4). Reconstitution of young male mice with aged female bone marrow enhanced cortical BV/TV as compared to age-matched female bone marrow, but did not alter cortical thickness (Figure 2A, B, D). No changes in trabecular BV/TV, trabecular number, or trabecular spacing were noted (Figure 2D-F), but we did observe an increase in trabecular thickness (Figure 2G). These results mirror those we observed when aged male bone marrow was transplanted into young male recipients.

Enhanced cortical bone mass could be due to increased bone formation and/or a reduction in bone resorption at the periosteal or endosteal surfaces, respectively. To determine how bone remodeling was altered within our chimeric mice we examined periosteal and endosteal cortical bone perimeter. Irradiation increased endosteal perimeter when comparing mice within group 1 to non-irradiated controls, supporting that irradiation increases resorption at the endosteal surface. In contrast, mice transplanted with aged bone marrow within group 2 exhibited decreased endosteal perimeter (Figure 2C). Likewise, transplantation of female bone marrow diminished endosteal perimeter (Figure 2C). These results suggest that transplantation of aged bone marrow reduced endosteal resorption of cortical bone or lead to increased endosteal formation.

### Subsets of myeloid cells from aged bone marrow associate with enhanced cortical bone

Our results from the bone marrow transplantation assays combined with recent work demonstrating that myeloid lineage cells (e.g., erythromyeloid progenitors, osteoclasts, osteomorphs, osteomacs) regulate the bone remodeling process, suggest the existence of aged myeloid cells that promote maintenance of cortical, but not trabecular bone. Moreover, a recent study demonstrated that erythromyeloid progenitors expand during intensive hematopoietic regeneration (20); therefore, we profiled myeloid cells within the bone marrow of our chimeric mice. We first evaluated the cellular subsets in young male recipients transplanted with sex-matched bone marrow derived from either age-matched or 40-week-old mice via singe cell RNA-sequencing (scRNA-Seq) of CD11B^+^ cells. Integration of these two datasets did not uncover unique clusters (Figure 3A), but we noted differential gene expression within clusters when comparing CD11B^+^ cells from mice reconstituted with aged as compared to age-matched bone marrow from either males (Figure 3) or females (Figure 4).

**Figure 3.**
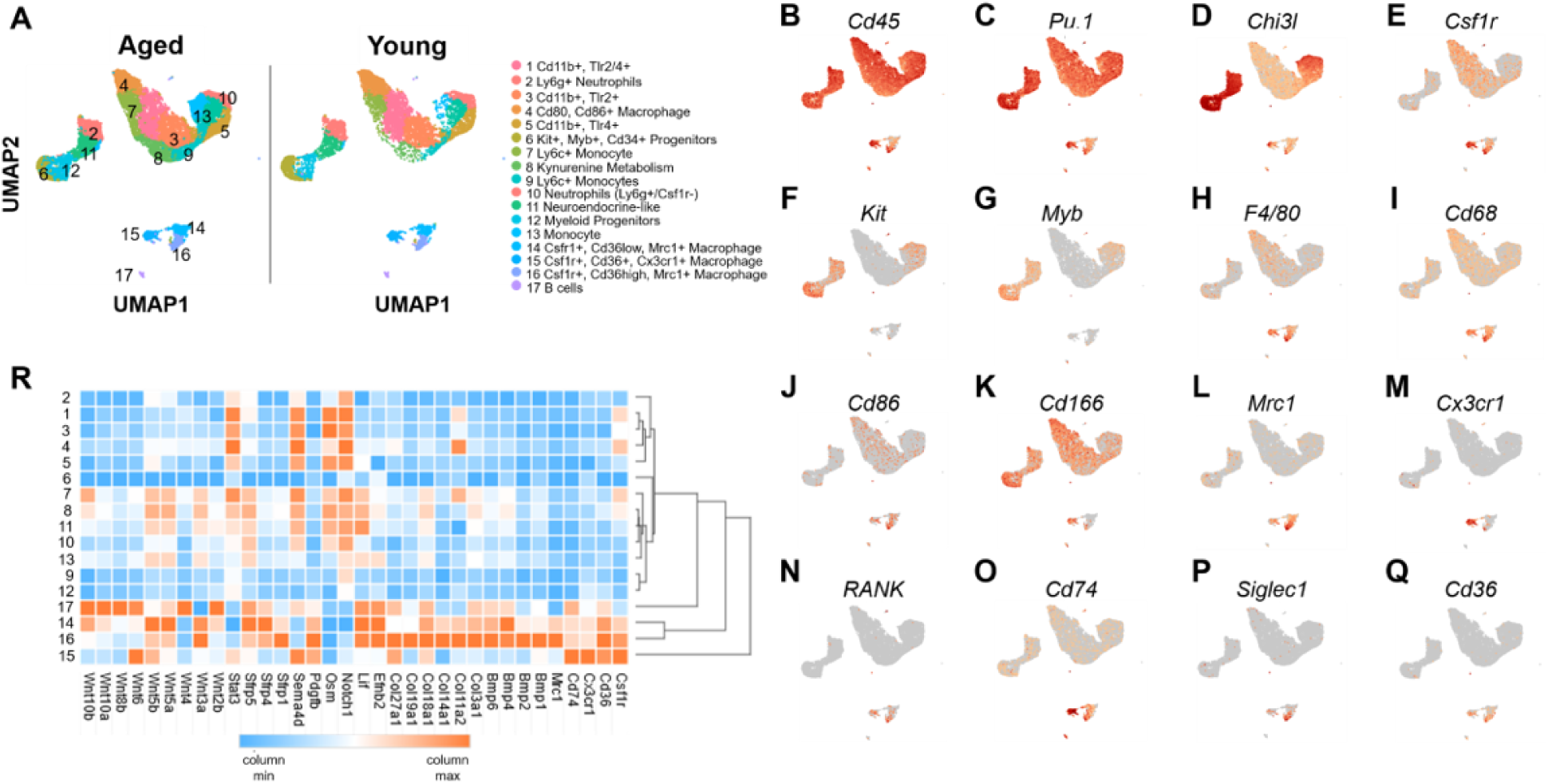
Identification of Anabolic Bone Marrow Subsets. (A-R) Young male recipient mice were lethally irradiated and reconstituted with T cell-depleted bone marrow (BM) from 1) young male or 2) aged male donor mice. Recipient mice were aged to 24-weeks for subsequent analyses. CD11B^+^ cells were collected from mice in groups 1 and 2 above and scRNA-Seq was performed using 10X Genomics technology. Resulting data were integrated and comparative analyses were performed. (A) UMAP 2D representation of CD11B^+^ clusters derived from recipient mice transplanted with aged and young bone marrow. (B-Q) Expression of surface markers used to differentiate clusters. (R) Heatmap of top differentially expressed genes from mice in group 2 vs 1 in clusters 14-16 across all clusters.

**Figure 4.**
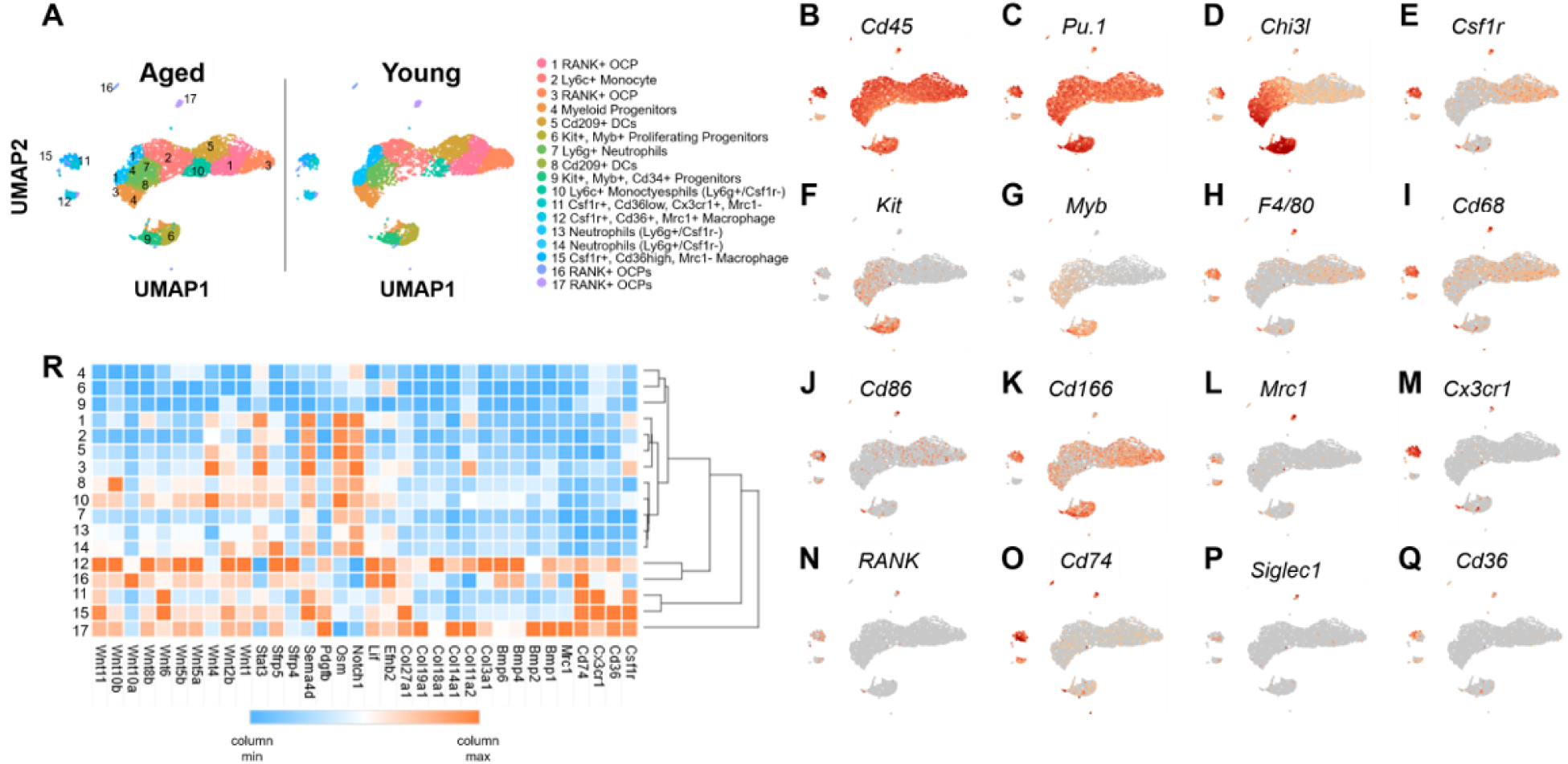
Identification of Anabolic Bone Marrow Subsets. (A-S) Young male recipient mice were lethally irradiated and reconstituted with T cell-depleted bone marrow (BM) from 1) young female or 2) aged female donor mice. Recipient mice were aged to 24-weeks for subsequent analyses. CD11B^+^ cells were collected from mice in groups 1 and 2 above and scRNA-Seq was performed using 10X Genomics technology. Resulting data were integrated and comparative analyses were performed. (A) UMAP 2D representation of CD11B^+^ clusters derived from recipient mice transplanted with aged and young bone marrow. (B-Q) Expression of surface markers used to differentiate clusters. (R) Heatmap of top differentially expressed genes from mice in group 2 vs 1 in clusters 14-16 across all clusters.

In mice reconstituted with male cells, all clusters expressed markers for hematopoietic cells, including *Cd45*, *Pu.1* and *Chi3l* (Figure 3B-D). We also noted multiple *Csf1r*^+^ clusters, including clusters 1, 4, 7, 8, 14, 15, and 16 (Figure 3E, R). Expression of *Kit* and *Myb* marked highly proliferative cells within clusters 6 and 12, which are likely myeloid progenitor cells (Figure 3F, G). Macrophage markers *F4/80*, *Cd68*, and *Cd86* were likewise expressed by cells within clusters 1, 4, 7, 8, and 14-16 (Figure 3H-J). The putative osteomac marker *Cd166* (21) was broadly expressed (Figure 3K), whereas the macrophage marker *Mrc1* was mostly restricted to clusters 14-16 (Figure 3L, R). Cells within cluster 16 expressed the highest levels of *Mrc1*, but expression was diminished by cells within cluster 15 (Figure 3L, R). Similarly, expression of *Cx3cr1* was restricted to clusters 14-16, with the highest expression observed by cells within cluster 15 (Figure 3M, R). Some cells within cluster 14-16 also expressed *RANK* (Figure 3N), supporting the potential of these cells to give rise to osteoclasts. We also noted high enrichment of *Cd74*, a putative osteomorph marker (22), *Siglec1/Cd169*, and *Cd36* within clusters 14-16 (Figure 3O-R). Recent genetic evidence suggests that cortical bone may be controlled specifically by several signaling pathways/components (e.g., Wnt and Notch signaling) (23). We next evaluated expression of these and related genes across all clusters and noted high expression of Bmps, Collagens, and Wnts by cells within clusters 14-16, and to some extent cluster 17 (Figure 2R). Enrichment of B Cell markers distinguished cluster 17.

We next evaluated the effect of sex on subsets of CD11B^+^ cells following intensive bone marrow reconstitution. We integrated scRNA-Seq results from the CD11B^+^ cells within the bone marrow of young male mice reconstituted with either age-matched male bone marrow or 40-week-old female bone marrow. This analysis did not reveal unique clusters (Figure 4A). As above, *Cd45*, *Pu.1*, and *Chi3l* were broadly distributed (Figure 4B-D). *Csf1r*^+^ clusters included clusters 1-3, 11, 12, 15 and 17 (Figure 4E). Myeloid progenitor cells marked by expression of *Kit* and *Myb* were contained within clusters 6 and 9 (Figure 4F, G). The cells within clusters 1-3, 5, 11, 12, 15, and 17 expressed the macrophage markers *F4/80*, *Cd68*, and *Cd86* (Figure 4H-J); *Cd166* was broadly expressed (Figure 4K). In contrast, *Mrc1* was expressed by cells within clusters 12 and 17 (Figure 4J, R). High expression of *Cx3cr1* marked cells within clusters 11 and 15, whereas expression was diminished with clusters 12 and 17 (Figure 4M, R). Expression of RANK was observed by cells within clusters 11 and 15-17 (Figure 4N). High expression of *Cd74* was also noted of clusters 11, 12 and 15-17 (Figure 4O, R), whereas *Cd36* was localized to cells within clusters 11, 12, 15 and 17, with cluster 11 exhibiting lower expression (Figure 4Q, R). As above, we noted enriched expression of Bmps, Collagens, and Wnts by cells within clusters 12, 15 and 17 (Figure 4R). We observed similar findings when comparing CD11B^+^ cells from young male mice reconstituted with bone marrow from 40-week-old males to those reconstituted with 40-week-old females, demonstrating that age, but not sex of donor cells associated with enhanced cortical bone mass. Likewise, our data suggest that specific subsets of CD11B^+^ cells express bone anabolic genes, and that these subsets are delineated by differential expression of *Cd36*.

### Characterization of CD11B^+^ cellular subsets within the bone marrow of young and aged animals

We noted the presence of several cellular subsets of cells within aged mice that expressed bone anabolic genes. To broadly define the types of myeloid cells within the bone marrow and evaluate how age alters these cellular subsets, we performed single cell RNA-sequencing (scRNA-Seq) on CD11B^+^ cells from the bone marrow of 8-or 40-week-old male and female C57Bl6/J mice. We chose these ages to reflect a period of bone anabolism (e.g., gain in bone mass) during youth and a period of age-associated bone loss prior to reproductive senescence. scRNA-Seq of CD11B^+^ cells from young animals revealed multiple cells types, including *RANK*^+^ osteoclast progenitors, dendritic cells, neutrophils, monocytes, macrophages as well as proliferative *Kit*^+^/*Myb*^+^ myeloid progenitors (Figure 5A, C).

**Figure 5.**
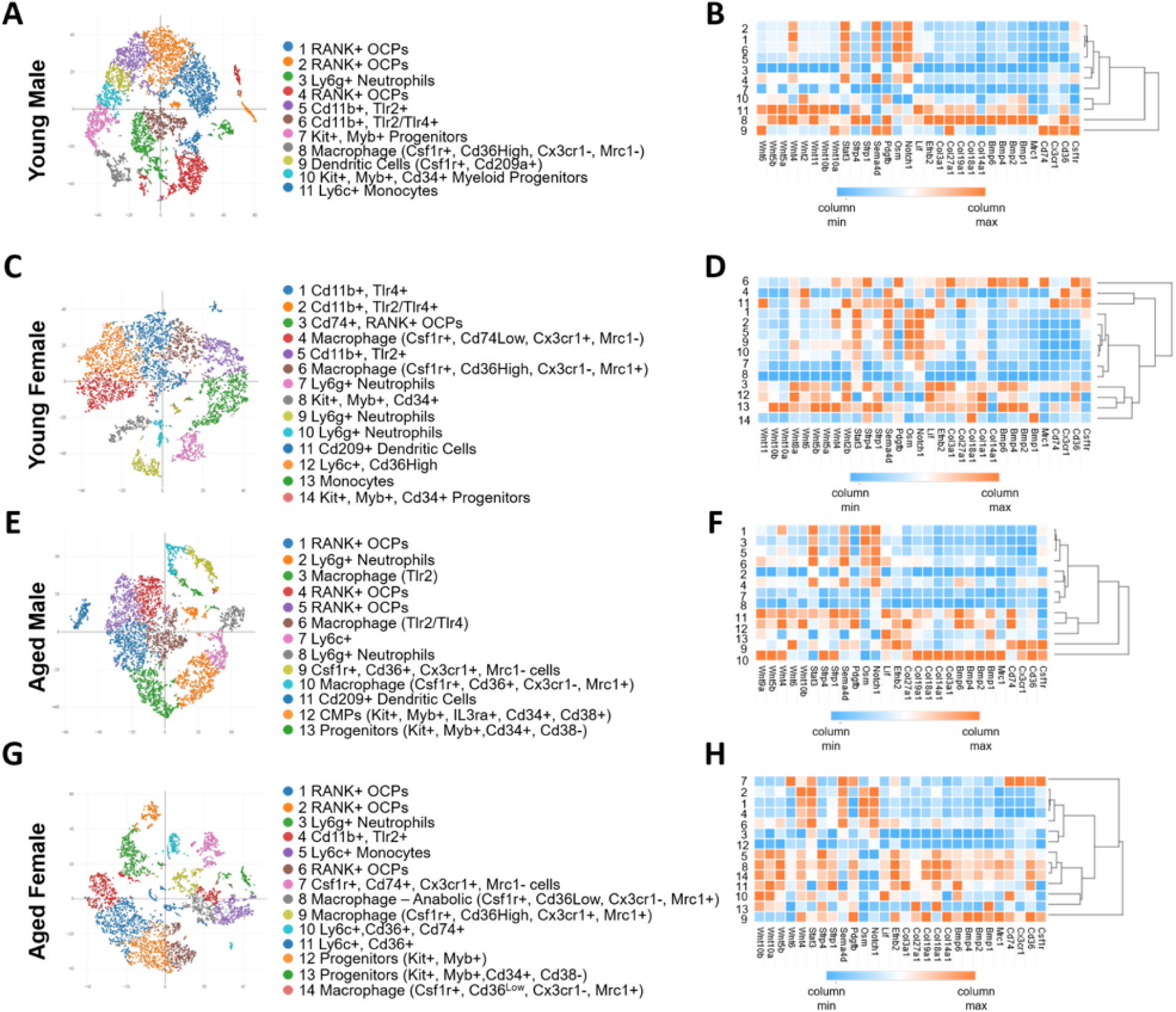
Single Cell RNA-Seq of CD11B^+^ Cells from Male and Female Bone Marrow Reveals Subsets of Aged Myeloid Cells that Express Anabolic Cytokines. (A-H) CD11B^+^ cells from the bone marrow of 8-or 40-week-old male and female mice were collected and single cell RNA-Seq was performed. (A) tSNE representation of data obtained from 8-week-old males. (B) Heatmap and hierarchical clustering of differentially expressed genes from clusters yielded from 8-week-old male data. (C) tSNE representation of data obtained from 8-week-old females. (D) Heatmap and hierarchical clustering of differentially expressed genes from clusters yielded from 8-week-old female data. (E) tSNE representation of data obtained from 40-week-old males. (F) Heatmap and hierarchical clustering of differentially expressed genes from clusters yielded from 40-week-old male data. (G) tSNE representation of data obtained from 40-week-old females. (H) Heatmap and hierarchical clustering of differentially expressed genes from clusters yielded from 40-week-old female data.

We first examined macrophage clusters within young males and females. We noted two macrophage clusters within 8-week-old CD11B-expressing bone marrow cells. Cluster 8 exhibited reduced expression of *Csf1r*, *Cx3cr1* and *Cd74*, but higher expression of *Mrc1* and *Cd36* (Figure 5B). Cluster 8 showed enrichment in bone anabolic genes such as BMPs, Collagens and Wnts. These bone anabolic proteins were also expressed by *Csf1r*^-^ cells within cluster 11. Likewise, we identified three *Csf1r*-expressing clusters within CD11B^+^ cells derived from the bone marrow of 8-week-old females (Clusters 3, 4, and 6, Figure 5C, D). Clusters 3 and 6 demonstrated reduced expression of *Csf1r*, but levels of *Mrc1* and *Cd36* were greater (Figure 5D). Cluster 4 demonstrated the highest enrichment for *Csf1r* and *Cx3cr1*, but exhibited little expression of *Cd36* or *Cd74*. Bone anabolic genes were enriched in clusters 3 and 6 (Figure 5D). These data pointed to a set of macrophages within the bone marrow that produce bone anabolic factors.

When examining aged CD11B^+^ cells, we likewise noted several myeloid subsets that were analogous to those present within young animals. Within males, we noted three subsets (clusters 4, 9, and 10) that all showed enrichment for *Csf1r* and *Cx3cr1* (Figure 5E, F). Moreover, clusters 9 and 10 demonstrated high expression of *Cd36*, but enrichment in *Cd74* by cells within cluster 9 and *Mrc1* by cells in cluster 10 differentiated these clusters (Figure 5F). All three clusters exhibited enrichment in Wnt gene expression, but only clusters 9 and 10 significantly expressed Bmps. Within CD11B^+^ female ages cells, clusters 7, 8, 9, and 14 all showed enrichment for *Csf1r*, *Mrc1* and *Cd36* (Figure 5G, H). *Cx3cr1* expression defined cluster 7 as well as cluster 9, but to a much lesser extent. *Cd74* also demonstrated differential expression amongst these four clusters (Figure 5H). Cluster 7 showed limited expression of Bmps and Wnts, whereas these bone anabolic genes were enriched amongst clusters 8, 9, and 14 (Figure 5H). These data highlight subsets of CD11B^+^/*Csf1r*^+^ cells within the bone marrow of aged animals that can be delineated by differential expression of *Cd74*, *Cx3cr1*, *Mrc1*, and/or *Cd36*.

### CD36+ cells associate with sites of bone resorption on endocortical surfaces

*Cd36* represented one of the most highly differentially expressed genes in our scRNA-Seq analyses and was exclusively expressed by specific subsets of cells within our analyses. We postulated that CD11B^+^CD36^+^ cells either serve as osteoclast progenitors or play a regulatory role in the bone remodeling process given that they are enriched in chimeric mice with elevated cortical bone mass. To investigate their localization and potential function, we performed immunohistochemical (IHC) staining for CD36 on femur sections from 12-and 24-week-old female mice. We also examined sections of cortical bone defects collected 14 days post-injury (24). To visualize bone remodeling sites, we additionally performed TRAP staining. Our results showed that CD36 was diffusely expressed by cells within the BM cavity, with no enrichment near cortical bone remodeling sites in 12-week-old female mice (Figure 6A, left column). However, in 24-week-old mice and within cortical bone defects, we observed strong CD36 staining in cells located near osteoclasts (Figure 6A, middle and right columns). CD36 expression by mature osteoclasts was not detected under any conditions. These results suggest that CD36^HI+^ cells may regulate bone remodeling and regeneration.

**Figure 6.**
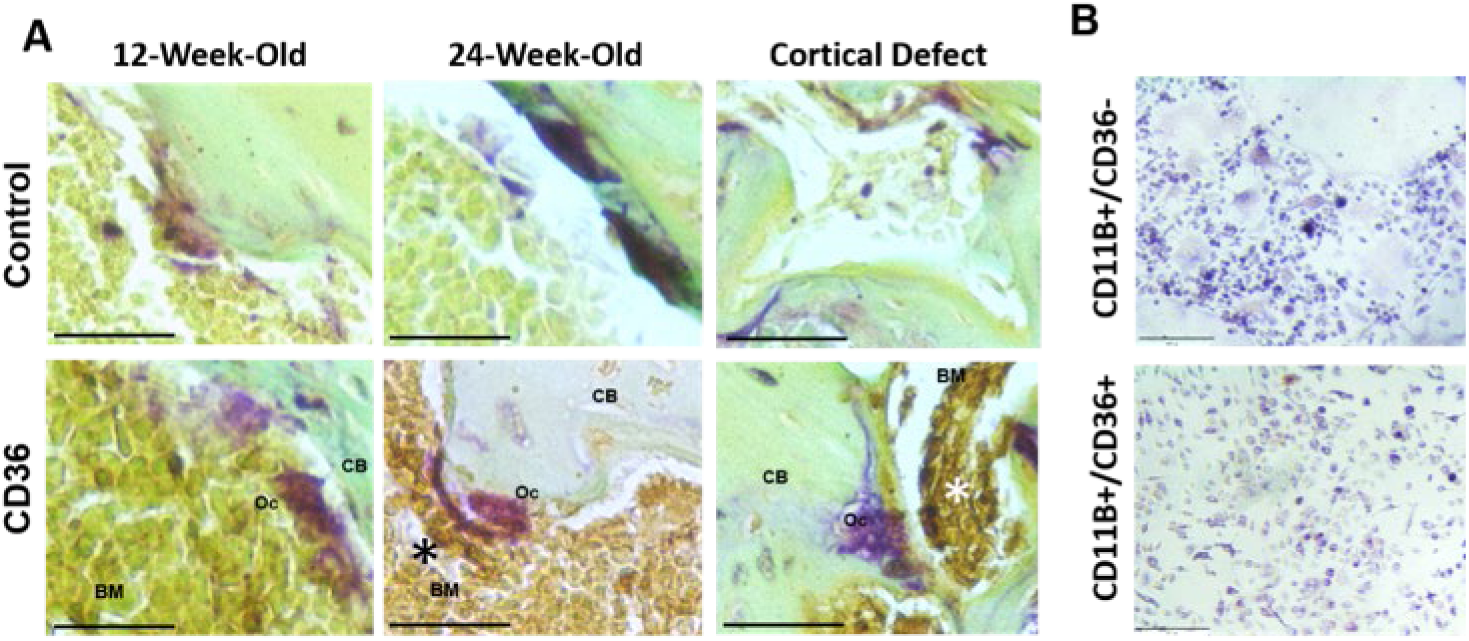
CD36^+^ Cells Associate with Sites of Cortical Bone Regeneration, but are Not Osteoclast Progenitors. (A) Immunohistochemical staining for CD36 of paraffin sections from uninjured 12-and 24-week-old females (left and middle panels, respectively) or cortical bone defects (right panels). Sections are counterstained with TRAP (purple) and fast green (n=5/group). BM: bone marrow, CB: cortical bone, OC: osteoclast (B) CD11B^+^CD36^-^ and CD11B^+^CD36^+^ BM cells were isolated and cultured in the presence of RANKL and M-CSF. TRAP staining was performed on day 4 (n=3).

To assess whether CD11B^+^CD36^+^ cells function as osteoclast progenitors, we isolated CD11B^+^CD36^+^ and CD11B^+^CD36^-^ cells from the BM of 6-9-week-old female mice and cultured them under osteoclastogenic conditions. While the CD11B^+^CD36^-^ population formed TRAP^+^ multinucleated cells, indicative of osteoclasts, the CD11B^+^CD36^+^ cultures only produced mononuclear cells (Figure 6B, lower panel). These findings suggest that CD11B^+^CD36^+^ cells are involved in cortical bone regeneration and remodeling but do not form osteoclasts, supporting previous reports (25).

### CD11B^+^CD36^+^ myeloid cells support bone formation

Our scRNA-Seq analysis revealed that CD11B^+^CD36^+^ cells express several bone anabolic factors, including *Wnt10b*, *Wnt6*, *Wnt3a*, *Bmp1*, and *Bmp2* (Figure 3R). To validate these findings, we FACS-sorted CD11B^+^CD36^+^ cells from the bone marrow of 6-9-week-old female mice. CD11B^+^CD36^+^ cells comprised 8.7% of the isolated bone marrow cells (Figure 7A). We observed increased expression of *Wnt10b*, *Wnt6*, and *Bmp1*, and reduced expression of *Wnt3a* and *Bmp2* by CD11B^+^CD36^+^ cells compared to CD11B^+^CD36^-^ cells (Figure 7B-F). *Wnt6* showed the most significant differential expression, which we further confirmed by western blotting (Figure 7G).

**Figure 7.**
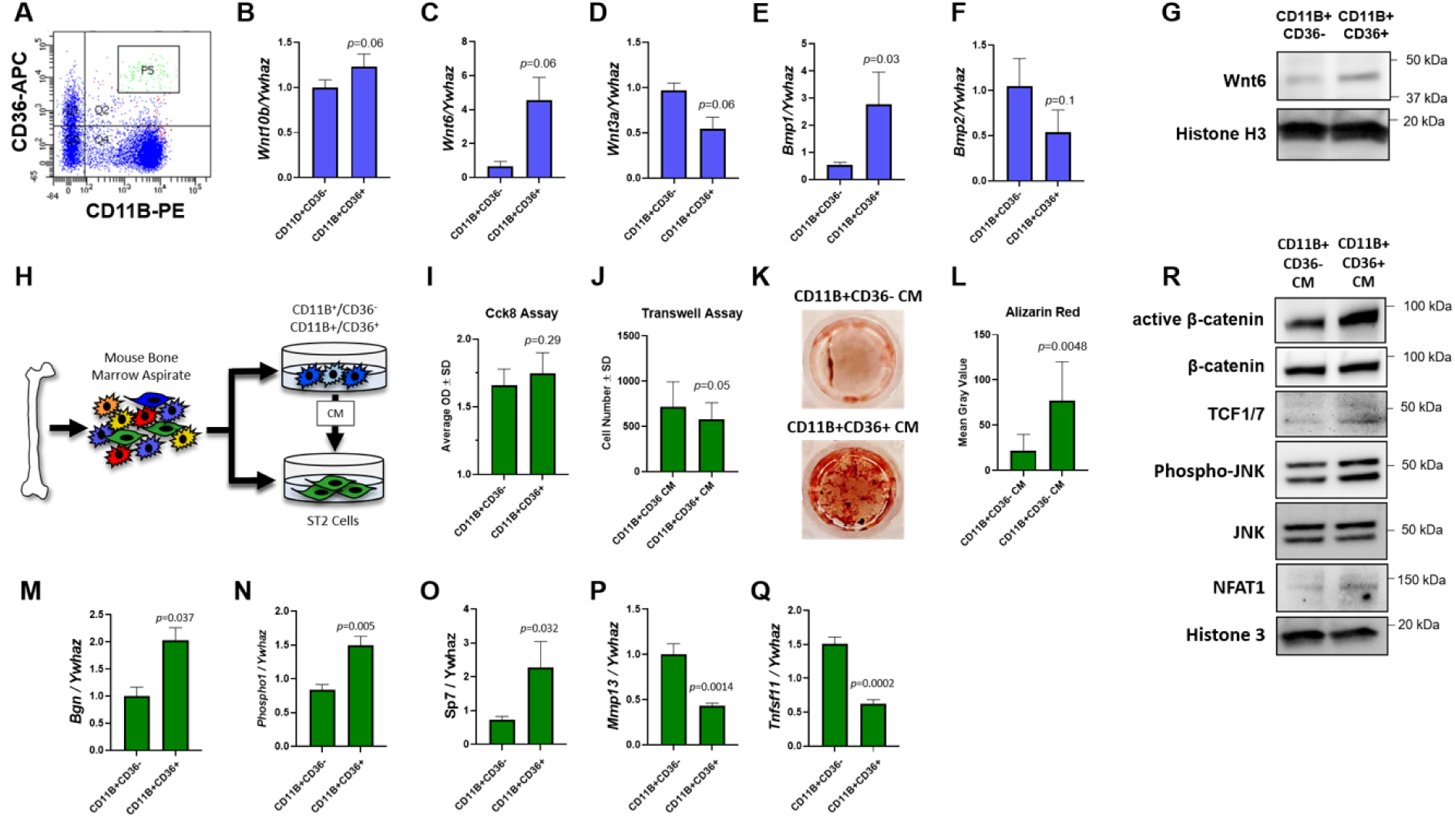
CD11B^+^CD36^+^ Cells Promote Bone Formation. (A) Flow cytometry. CD11b+CD36-and CD11b+CD36+ cells were sorted and gene expression was evaluated by qPCR (B-F) and western blotting (G), n=3, *p* values as shown. (H) Experimental design for (I-Q) Osteoprogenitor cells were treated with CM for 24 hours and (I) Cck8 and (J) transwell assays were performed. Osteoprogenitor cells were treated with CM for 8 weeks and Alizarin Red staining (K) was performed and quantified (L), n=3, p values as shown. RNA was collected and gene expression (M-Q) was assessed by qPCR, n=3, p values as shown. (R) Osteoprogenitor cells were treated with CM for 60 minutes and western blotting was performed (n=3).

Next, we investigated whether factors secreted by CD11B^+^CD36^+^ cells influence osteoprogenitor proliferation, migration, commitment, and mineralization. CD11B^+^CD36^-^ and CD11B^+^CD36^+^ cells were sorted from the BM of 6-9-week-old male and female C57Bl/J mice, cultured with M-CSF for three days, and conditioned medium was collected. ST2 osteoprogenitor cells were then cultured in conditioned medium for 24 hours to assess proliferation and migration (Figure 7H). Cck-8 assays indicated no significant impact on osteoprogenitor proliferation and transwell assays similarly showed no significant effects on migration. To evaluate osteoprogenitor commitment and mineralization, ST2 cells were cultured in conditioned medium under osteogenic conditions. Enhanced mineralization was observed in cultures treated with CD11B^+^CD36^+^ conditioned medium (Figure 7K, L) accompanied by increased expression of *Bgn*, *Phospho1*, and *Sp7* (Figure 7M-O). Conversely, *Mmp13* and *Tnfrsf11*/RANKL expression was reduced in these cultures (Figure 7P, Q). Finally, we assessed the activation of β-catenin and JNK signaling pathways in ST2 cells cultured in conditioned medium from CD11B^+^CD36^-^ and CD11B^+^CD36^+^ cells for one hour. Western blotting revealed elevated levels of active β-catenin and JNK phosphorylation, indicating increased canonical and/or non-canonical Wnt signaling (Figure 7R). Likewise, NFAT1 and TCF1/7 levels were enhanced (Figure 7R). These results suggest that CD11B^+^CD36^+^ cells produce WNT6, which promotes osteoblast differentiation and bone formation.

### CD11B^+^CD36^+^ Cells within Human Bone Marrow

To show that our results may translate to humans, we performed single cell sequencing of CD11B^+^ cells from the bone marrow aspirate of a 45-year-old human female. Using the pipeline described above, we aligned data to the hg38 reference genome. We identified a cluster of isolated CD11B^+^ cells exquisitely defined by *Cd36* (Figure 8A, Q, R). CD11B^+^CD36^+^ cells within cluster 7 also expressed *Cd45*, *Pu.1*, *Csf1r*, *F4/80*, *Cd68*, *Cd86*, *Cd166*, *Cd74*, and *Cx3cr1*, but were mostly negative for *Chi31l*, *Mrc1*, *RANK*, and *Siglec1/Cd169* (Figure 8B-P). We also noted that expression of *Cd166*, the putative osteomac marker, and *Cd74* were broadly expressed by CD11B^+^ cells (Figure 8K, O). Expression of *Wnt6* was likewise restricted to CD11B^+^ cells cluster 7; thus, bone marrow from aged humans also contains an anabolic macrophage population marked by CD11B/CD36.

**Figure 8.**
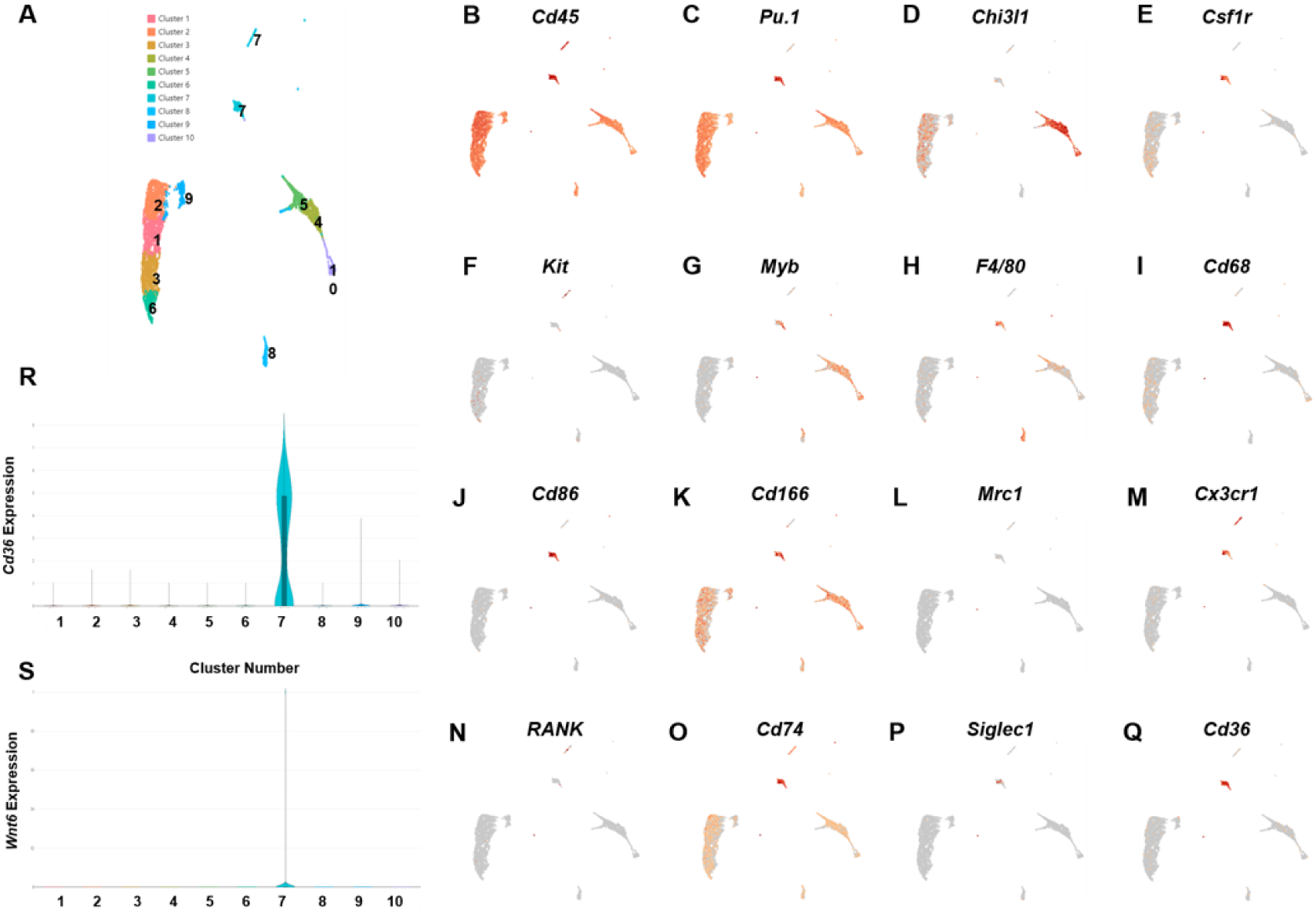
Identification of Human Bone Marrow Macrophage Subsets. CD11B^+^ cells were collected from the BM aspirate of a 45-year-old human female and scRNA-Seq was performed using 10X Genomics technology. (A) UMAP 2D representation of CD11B^+^ clusters. (B-Q) Expression of surface markers used to differentiate clusters. Violin plots of CD36 (R) and Wnt6 (S) expression.

## DISUCSSION

The goal of our study was to examine how age and sex of hematopoietic cells affect skeletal aging. Prior work documented that altered mesenchymal cell function facilitates age-associated changes to trabecular bone, with consequent myeloid cell skewing (26, 27). Our study supports this finding, as we did not observe changes to trabecular bone mass or microarchitecture. In contrast, our data support that age alters bone marrow composition following intensive hematopoietic regeneration as previously described (20), and those alterations in bone marrow composition facilitate gains in cortical bone. Moreover, our data suggest that a subpopulation of aged macrophages limit endocortical bone resorption.

Bone resorbing osteoclasts can arise from hematopoietic stem cells or from long-lived erythro-myeloid progenitors within the bone marrow. Osteoclasts have an estimated 6-month half-life in mice (28–30). Recent evidence suggests that osteoclasts derived from different subsets of progenitors may facilitate bone remodeling in different contexts (31–33). For instance, osteoclast progenitors that contribute to homeostatic bone remodeling differ from those mediating bone loss associated with inflammatory conditions (32). Osteoclasts that remodel bone in other perturbed states or within different bone compartments may likewise differ. Our data suggest that age-associated resorption at endosteal cortical bone surfaces may be distinct and limited by secreted factors produced CD11B^+^CD36^+^ cells. To the best of our knowledge, this represents the first example of an aged myeloid cell type that limits bone resorption.

In our study, we identified a specific myeloid subset of aged cells that limit age-associated cortical bone loss. Surface marker expression of CD11B and CD36 characterizes this subset of cells. In addition to these surface markers, osteomorph markers (e.g., Axl, Cadm1, Ccr3, Cd74, Vcam1 (22)) were significantly enriched within our identified myeloid subset. Osteomorphs have been speculated to function during the reversal phase of bone remodeling. Given their proximity to bone resorbing osteoclasts along the cortical surface, our identified cellular subset could be osteomorphs. Indeed, our scRNA-Seq analyses demonstrated expression of RANK within these clusters, but we were unable to generate osteoclasts from sorted CD11B^+^CD36^+^ cells in vitro. In contrast, CD11B^+^CD36^-^ cells efficiently formed osteoclasts in vitro. These data suggest that our identified myeloid subset is potentially distinct from osteomorphs, but we do not know if CD11B^+^CD36^+^ cells could fuse with existing osteoclasts in vivo.

CD36 is a cell surface marker delineating our population of interest, but we are unsure if CD36 is merely a marker for this myeloid subset, or if CD36 plays a functional role in controlling age-associated cortical bone loss. CD36 is a membrane glycoprotein that functions as a scavenger receptor for various ligands, including collagen, thrombospondin-1, anionic phospholipids, and long chain fatty acids. While CD36 is broadly expressed, adipocytes, cardiac myocytes, and immune cells including myeloid cell types (e.g., monocytes, macrophages) exhibit biased expression. CD36 mediates the formation of giant cells, but does not facilitate osteoclast fusion (25). Our data further confirm that CD11B^+^CD36^+^ cells do not form osteoclasts.

Global deletion of *Cd36* (i.e., *Cd36^-/-^*) resulted in conflicting effects on bone mass. In the first published study, male and female 1-to 6-month-old *Cd36^-/-^* mice demonstrated reduced vertebral and femoral cortical bone mass characterized by diminished bone formation (34). No changes in markers of resorption were noted (34). Furthermore, in vitro osteoblastogenesis from bone marrow stromal cells was diminished, though the impact of myeloid cell *Cd36* deletion in these cultures could be a confounding factor. Conversely, a later study reported increased vertebral and femoral cortical BV/TV in female 12-week-old *Cd36^-/-^* mice, with no detection of CD36 expression by mature osteoclasts or osteoblasts (35). Given these conflicting results using *Cd36* global knockout mice, an examination of the role of *Cd36* specifically within monocyte/macrophage-lineage cells during bone remodeling and regeneration is warranted as understanding the role of *Cd36* in bone biology would offer potential therapeutic targets for managing bone-related diseases and improving skeletal health.

Our scRNA-Seq data demonstrated enrichment of secreted factors (e.g., Wnts, Bmps) by aged myeloid cells within our clusters of interest. Coupled with decreased endosteal perimeter, we speculated that CD11B^+^CD36^+^ cells may limit resorption at endosteal surfaces. Wnt signaling limits osteoclastogenesis (36). Likewise, canonical Wnt signaling promotes expression of OPG by osteoblasts, leading to diminished osteoclastogenesis (37). In contrast, little is known about production of Wnt ligands by other cell types involved in bone remodeling and how this impacts cortical bone resorption. In fact, studies suggest that osteoclasts and bone forming osteoblasts may not be present at sites of bone remodeling at the same time (38–40), particularly at the endosteal surface. Likewise, age disrupts the temporal and spatial processes of resorption and formation. Our studies point to CD11D^+^CD36^+^ cells as targets to limit age-associated cortical bone loss. In future studies, we will determine factors produced CD11B^+^ CD36^+^ cells that limit age-associated osteoclastogenesis and bone resorption.

Our study has several limitations. We did not assess possible contributions of splenic cells to our phenotype. Studies have shown that osteoclast progenitors from the spleen can migrate to bone and participate in bone remodeling and repair (28, 31, 33). We did not observe significant expression of X-linked gene within the bone marrow of males reconstituted with female bone marrow or splenomegaly; nonetheless, we cannot completely rule out the contribution of these cells within our study. Although we speculate that CD11B^+^CD36^+^ cells are not osteomorphs based on their inability to support osteoclastogenesis in vitro, we do not know if this is true in vivo. Our definition of aged mice does not strictly fit the well-accepted definition (e.g., older than one year). We wanted to uncover mechanisms of age-associated bone loss independent of reproductive status to reflect what may be initiating events in the aging process as both men and women begin to lose bone mass starting in the third decade of life.

## MATERIALS AND METHODS

### Animal procurement and housing

The Jackson Laboratory provided male and female C57Bl6/J mice (strain #000664, Bar Harbor, ME, USA) aged to either 6-or 38-weeks old at time of purchase. Mice were acclimated to housing conditions for two weeks prior to experimentation. All animal research was conducted according to guidelines provided by the National Institutes of Health and the Institute of Laboratory Animal Resources, National Research Council. The University of Minnesota Institutional Animal Care and Use Committee approved all animal studies.

### T Cell Depletion of Whole Bone Marrow

Male and female 8-or 40-week-old mice were euthanized by CO_2_ asphyxiation. The femora and tibiae were collected and connective tissue was removed. Bone epiphyses were removed and the marrow cavity was flushed with sterile PBS. Whole bone marrow suspensions from age-matched mice were pooled and pelleted at 1000 rpm for five minutes. Cell pellets were then suspended in 1X RBC lysis buffer (#00-4333-57, ThermoFisher Scietific, Waltham, MA, USA) and incubated for five minutes. Cells were pelleted at 1000 rpm and suspended in MACS buffer (0.5% bovine serum albumin, 2mM EDTA, phosphate buffered saline) by diluting MACS BSA stock solution (Miletnyi Biotech, Bergisch Gladbach, North Rhine-Westphalia, Germany, #130-091-376) 1:20 with autoMACS rinsing solution (Miltenyi Biotech, #130-091-222,) to a density of 0.2x10^6^ cells/μl. Cell suspensions were then incubated with 1 μl PE-conjugated anti-CD8 and anti-CD3 antibodies per 1x10^7^ nucleated cells for 20 minutes in the dark at 4C. Cells were then washed with MACS buffer, pelleted and suspended at a density of 1.25x10^6^ cells/μl in MACS buffer. Anti-PE microbeads (20 μl/10x10^6^ cells, Miltenyi Biotech, #130-048-801) were added and cells were incubated at 4C for 20 minutes in the dark. Cells were then washed pelleted and applied to an LS column (#130-042-401, Miltenyi Biotech) placed in a MidiMACS Separator magnet (Miltenyi Biotec, #130-042-301h). Eluted CD8^-^/CD3^-^ bone marrow cells were pelleted and suspended in PBS at a density of 50x10^6^ cells/ml.

### Irradiation and bone marrow reconstitution

Six-week-old (e.g., young) male mice (n=34) were lethally irradiated using and X-ray irradiator via two successive does of 500 rads separated by a period of four hours. Two hours after completion of irradiation, mice were reconstituted via a 200 μl tail vein injection of 10x10^6^ T cell-depleted whole bone marrow cells suspended in PBS derived from either 6-week-old males (n=10), 6-week-old females (n=8), 40-week-old males (n=8), or 40-week-old females (n=8). Following bone marrow reconstitution, mice were housed in autoclaved cages and were administered 15 μg polymixin B sulfate and 40 μg/ml neomycin sulfate in their drinking water for two weeks. Mice were aged an additional 16 weeks (e.g., 24-weeks-old) and then euthanized for analyses.

### Micro-computed tomography

Right femora collected from 24-week-old chimeric mice and non-irradiated 24-week old males were fixed in 10% neutral buffered formalin for 48 hours, wrapped in gauze, and placed in 1.5mL screw-cap tubes filled with 70% ethanol. Scanning was performed using the XT H 225 micro-computed tomography (micro-CT) machine (Nikon Metrology Inc., Brighton, MI, USA) set to 120kV, 61µA, 720 projections at two frames per projection with an integration time of 708 milliseconds as previously described (41). Scans were done at an isometric voxel size of 7.11 µm with a 1mm aluminum filter, 17 minutes per scan. Each scan volume was reconstructed using CT Pro 3D (Nikon metrology, Inc., Brighton, MI, USA). Reconstructions were converted to bitmap datasets using VGStudio MAX 3.2 (Volume Graphics GmbH, Heidelberg, Germany). Scans were reoriented via DataViewer (SkyScan, Bruker micro-CT, Belgium) to create a new bitmap dataset for consistent analysis. From these data, representative transverse slices were selected for qualitative analysis. Morphometric analysis was performed using SkyScan CT-Analyzer (CTAn, Bruker micro-CT). Bruker’s instructions and guidelines for analysis within the field was followed throughout analysis (41). 3D Analysis of trabecular bone was performed in the distal metaphysis 0.7 mm proximal to the growth plate and extending 1.5 mm proximally toward the bone diaphysis. For the bone cortex, 2D analysis occurred in a 0.5 mm section within the mid-diaphysis defined as 4 mm from the growth plate. Regions of interest were set by automated contouring for the trabecular and cortical ranges, with some manual editing when necessary. Binary selection of all samples resulted in two separate global thresholds used to separate bone from surrounding tissue within the trabecular and cortical regions of interest. Parameters measured for trabecular bone include bone volume per total volume (BV/TV), bone surface per total volume (BS/TV), bone surface per bone volume (BS/BV) trabecular thickness, number, and spacing (Tb.Th, Tb.N, Tb.Sp), and connective density (Conn.D). Cortical parameters measured were BV/TV, periosteal and endosteal perimeter (pM), as well as cortical thickness (Ct.Th).

### CD11B^+^ magnetic assisted cell sorting

Bone marrow cells were flushed from the femora and tibiae of 6-month-old chimeric mice from each group or from 8-and 40-week-old male and female C57Bl6/J mice. Whole bone marrow suspensions resulting from each mouse were pelleted at 1000 rpm for five minutes. Cell pellets were then suspended in 1X RBC lysis buffer and incubated for five minutes. Cells were pelleted at 1000 rpm and suspended in MACS buffer to a density of 0.11x10^6^ cells/μl. Nucleated cells were then incubated with 10 μl CD11B microbeads (Miltenyi Biotech, #130-126-725) per 1x10^7^ cells for 10 minutes at 4C. Cell suspensions were then brought to a total volume of 500 μl with MACS buffer and applied to an equilibrated LS column placed in a MidiMACS Separator magnet. Columns were then washed three times with MACS buffer and removed from the magnet. CD11B^+^ cells were eluted from the column with 5 ml MACS buffer. Cells were then enumerated and used for subsequent scRNA-Seq analyses.

### Single cell RNA-sequencing

CD11B+ cells from chimeric mice in each group or 8-and 40-week-old male and female C57Bl6/J mice were isolated via MACS as described above. Single cell cDNA libraries from each sample were the prepared using Chromium Single cell 3′ Reagent v3 Kits (10× Genomics, Pleasanton, CA, USA) according to the manufacturer’s protocol. Each single-cell suspension was mixed with primers, enzymes, and gel beads containing barcode information and then loaded on a Chromium Single Cell Controller (10× Genomics) to generate single-cell gel beads in emulsions (GEMs). Each gel bead was bonded to one cell and encased within oil surfactant. After generating the GEMs, reverse transcription was performed using barcoded full-length cDNA followed by the disruption of emulsions using the recovery agent and cDNA clean up with DynaBeads MyOne Silane Beads (ThermoFisher Scientific, #37002D). cDNA was then amplified by PCR with an appropriate number of cycles and thermal conditions that depended on the recovery cells. Subsequently, the amplified cDNA was fragmented, end-repaired, A-tailed, ligated to an index adaptor, and subjected to library amplification. Quality of amplified cDNA was assessed using an Agilent Bioanalyzer prior to high-throughput sequencing. Following quality control, the cDNA libraries were sequenced on an Illumina NovaSeq 6000 sequencer. The Genomics Core at the University of Minnesota performed library construction and sequencing.

### Single Cell Bioinformatic Analyses

10× Genomics Cell Ranger software (version 6.0.0) was used to process the raw scRNA-seq data in accordance with the guidelines specified by the manufacturer. Cell Ranger software (2020-A, 10× Genomics Cell Ranger Count v7.0.1) was used to demultiplex the raw base call (BCL) files produced by the Illumina NovaSeq 6000 sequencer. Next, Cell Ranger was used to count the unique molecular identifiers (UMI) and barcodes and align fastq files to the mouse reference genome (GRCm39). Quality assurance and subsequent studies were then performed on the feature-barcode matrices created by Cell Ranger. The study was filtered to exclude cells with outlier status, aberrant gene detection rates (<500 and >5,000), and high mitochondrial transcript level (>8%, a sign of cellular stress). Cells were then organized into an ideal number of clusters for de novo cell type discovery. Then, using uniform manifold approximation and projection (UMAP), a non-linear dimensional reduction was carried out, enabling for the identification and visualization of different cell clusters. Data integration was performed to compare between samples.

### CD11B^+^CD36^+^ Cell Isolation

Whole bone marrow was flushed from the hind limbs of 6-8-week-old male and female C57Bl6/J mice (n=3 per sex) and red blood cells were lysed with 1X RBC lysis buffer as described above. Nucleated cells from whole bone marrow were pelleted at 1000 rpm for 5 minutes, and the supernatant was removed and discarded. Nucleated cells were divided into two subsets, one for isolation of all CD11B^+^ cells within the bone marrow and the other for isolation of CD11B^+^/CD36^-^ and CD11B^+^/CD36^+^ cells. Cells were counted and pelleted at 1000 rpm. Nucleated cells within the pellet were suspended in 100 μl MACS Buffer per 1x10^7^. For isolation of CD36^+^ cells, cells were then incubated with Allophycocyanin (APC)-conjugated anti-CD36 (10 μl per 1x10^7^ cells, Miltenyi Biotech, #130-122-085) antibodies for 10 minutes at 4C. Cells were pelleted, washed, suspended in 1X MACS buffer (80 μl per 1x10^7^ cells) and incubated with anti-APC microbeads (20 μl per 1x10^7^ cells, Miltenyi Biotech, #130-090-855) for 15 minutes at 4C. Cells were washed and then suspended in 500 μl MACS buffer. An LS MACS separation column was placed within the magnetic field of a MACS Separator magnet and the column was equilibrated with 3 ml MACS buffer. Following equilibration, anti-APC microbead-labeled cells were applied to the LS column. The column was washed 3X with 3 ml MACS buffer. Unbound cells were used to isolate CD11B^+^CD36^-^ cells. To isolate APC-labeled cells, the column was removed from the magnet and cells were flushed from the column with 5 ml 1X MACS buffer. Eluted cells were then pelleted at 1000 rpm for 5 minutes, and the supernatant was removed and discarded. To isolate CD11B^+^ cells from each of the three fractions (e.g., whole bone marrow, CD36^-^ cells, and CD36^+^ cells), cells were then suspended in 100 μl 1X MACS Buffer per 1x10^7^ cells and incubated with phycoerythrin (PE)-conjugated anti-CD11B (10 μl per 1x10^7^ cells, Miltenyi Biotech, #130-113-235) for 10 minutes at 4C. Cells were washed, suspended in MACS buffer (80 μl per 1x10^7^ cells), and incubated with anti-PE microbeads (20 μl per 1x10^7^ cells, Miltenyi Biotech, #130-048-801) for 15 minutes at 4C. Magnetic sorting for PE-labeled cells was accomplished as described above. To validate effective sorting of CD36^+^CD11B^+^ cells via MACS, flow cytometry was performed (n=3). For flow cytometry analysis, we prepared single stained controls and double stained sample with unsorted nucleated cells with PE-conjugated anti-CD11B, APC-conjugated anti-CD36, or co-stained with anti-CD11B and anti-CD36 antibodies (1x10^6^ cells per condition), respectively. Data were acquired on a BD LSR II (H1010) and analyses were performed using BD FACSDiva^TM^ software.

### Culture of CD36^+^CD11B^+^ cells

MACS sorted CD11B^+^, CD11B^+^/CD36^-^ and CD36^+^/CD11B^+^ cells were isolated via MACS and cultured at a density of 0.26x10^6^ per cm^2^ in α-Minimal Essential Medium supplemented with 50 ng/ml recombinant mouse macrophage-colony stimulating factor (M-CSF) (#416-ML, R&D Systems, Minneapolis, MN, USA). Cells were maintained in culture and supplemented with M-CSF for 7 days or until adherent. Medium conditioned by each cell fraction was then collected, filtered with a 70 μm filter to remove cellular debris, and stored at -80C before subsequent analyses. Cells were lysed in TriZol reagent (ThermoFisher Scientific, #15596018) for gene expression analyses. For osteoclastogenesis experiments, cells from each fraction were plated at a density of 0.4x10^6^/cm^2^ in αMEM supplemented with 100 ng/ml RANKL (R&D Systems, #462-TR-010) and 50 ng/ml M-CSF. Cells were refed with αMEM supplemented with RANKL and M-CSF every other day for seven days, fixed with 10% neutral buffered formalin, and TRAP stained using the Leukocyte Acid Phosphatase (TRAP) kit (Sigma-Aldrich, #387A-1KT).

### RNA Isolation and Semi-quantitative RT-PCR

Total RNA was extracted from human macrophages using TRIzol and chloroform. Reverse transcription using the iScript™ Reverse Transcription Supermix (Biorad, #1708841) using 1 μg of total RNA was used to generate cDNAs. Gene expression was then assessed via real-time PCR using gene-specific primers listed in Table 1. Fold changes in gene expression for each sample were calculated using the 2^−ΔΔCt^ method relative to control after normalization of gene-specific C_t_ values to Ywhaz C_t_ values (24, 42). Shown are averaged data from three independent replicate experiments derived from female cultured cells.

**Table 1.**
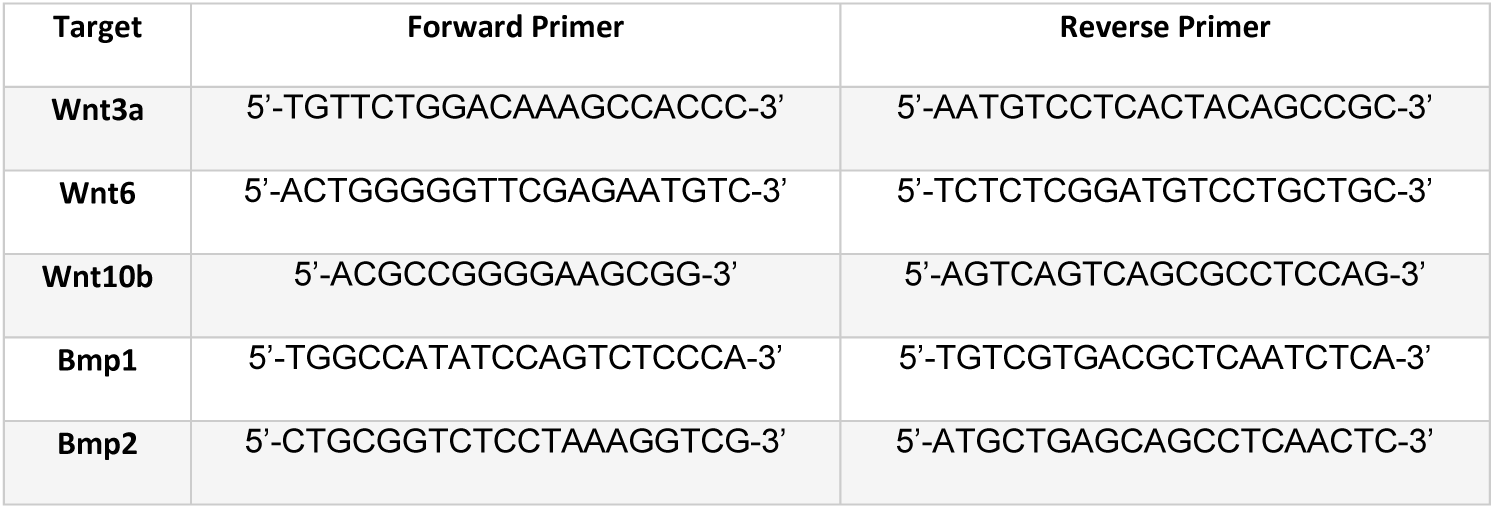
Gene specific primers used for semi-quantitative RT-PCR.

### Cortical Bone Defect Generation

Single-cortex, fully stabilized, mid-diaphysis femoral defects were generated in the right femur as previously described (24, 43). Mice in all experimental groups were given Buprenorphine SR provided perioperatively at 0.5 mg/kg given 2-4 hours before surgery via subcutaneous injection. Mice were then anesthetized with isoflurane and prepared for aseptic surgery. A small incision was made in the skin overlying the right hamstring and the femoral bone shaft was exposed by blunt dissection of the underlying muscle. A 0.7 mm diameter steel burr drill bit (#19007-07, Fine Science Tools, Foster City, CA) and an electric drill was used to induce a single-cortex defect in the mid-diaphysis of the femur. Defects were immediately irrigated with 1 ml sterile saline followed by incision closure. Mice were sacrificed by carbon dioxide inhalation on postoperative day 14.

### Immunohistochemistry (IHC) and TRAP Staining

Tibiae from 24-week-old female mice were collected, fixed in 10% neutral buffered formalin for 24 hours and then stored in 70% ethanol. Following decalcification in 15% EDTA for 14 days, tissues were paraffin embedded and 7 micron sections were collected. IHC staining was performed with antibodies directed to CD36 (#HPA002018, Sigma-Aldrich), or an irrelevant control. Detection was accomplished using the Mouse and Rabbit Specific HRP/DAB (ABC) Detection IHC Kit (#ab64264, Abcam, Cabmbridge, UK). Sections were the TRAP stained and then counterstained with Fast Green as previously described (42).

### Measurements of proliferation and osteoblastogenesis

ST2 cells were cultured at a density of XXX cells/cm2 in CD11B^+^, CD11B^+^/CD36^-^ or CD11B^+^/CD36^+^ conditioned medium for 24 hours. MSC proliferation was then determined using a Cck8 assay (Cell Counting Kit, Sigma Aldrich, #96992-100TESTS-F). To assess MSC migration, transwell assays were performed using Corning Costar transwell assay cell culture inserts (Sigma Aldrich, #CLS3464-12EA). Before starting the assay, the transwell were equilibrated over night with MEM (no FBS). Next day, ST2 cells were trypsinized and collected in 0.2% FBS containing MEM. A total of 2x10^4^ cells were seeded into the upper chamber of the transwell inserts. The lower chamber was filled with an equal mixture of conditioned medium and MEM (10% FBS). Then, the plate was incubated for 18hrs to allow migration. Following incubation, the plate was taken out and the media was removed. The insert was washed with 1x PBS two times. The non-migrated cells were carefully removed from the upper side of the membrane using cotton swab and the migrated cells on the other side of the membrane was fixed with 100% methanol, for 5 min at room temperature and stained with Giemsa stain for 10min. Excess stain was removed by washing the chamber with distilled water and then air drying The membrane was carefully cut out using a scalpel blade and placed on slide. The membrane was mounted on the slide using VECTASHIELD® Antifade Mounting Medium with DAPI. Cells were visualized under the fluorescent microscope, and five random fields per insert were counted using Image J.

### Western Blotting

Cell lysates were collected in a buffered SDS solution (0.1% glycerol, 0.01% SDS, 0.1 m Tris, pH 6.8) on ice. Total protein concentrations were obtained with the Bio-Rad D_C_ assay (Bio-Rad). Proteins (20 μg) were then resolved by SDS-PAGE and transferred to a polyvinylidene difluoride membrane. Western blotting was performed with antibodies (1:2000 dilution) for WNT6 (Thermo Fisher Scientific, #24201-1-AP), Active β-catenin (Millipore, #05-665-25UG), β-catenin (BD Biosciences, #610153), TCF1/7 (Cell Signaling Technology, 2203T), phospho-JNK (Cell Signaling Technology, #4668T), JNK (Cell Signaling Technology, #9252T), NFAT1 (Cell Signaling Technology, #4389S), Histone 3 (Millipore, #05-928), and corresponding secondary antibodies conjugated to horseradish peroxidase (HRP) (Cell Signaling Technology). Antibody binding was detected with the Supersignal West Femto Chemiluminescent Substrate (Pierce Biotechnology, Rockford, IL). Shown are data from averaged from three males, but studies were performed in females (n=3 per group) in three independent replicate experiments, each containing six pooled replicate wells per group.

## ACKNOWLEDGEMENTS

The authors would like to thank the support from the Comparative Pathology Shared Resource, the Minnesota Dental Research Center for Biomaterials and Biomechanics, and technical assistance from Mr. David HH Molstad. This work was made possible by research grants from the National Institute for Musculoskeletal, Arthritis, and Skin Diseases (R21AR084530), a seed grant from the UMN BIRCWH award (HD055887), and the University of Minnesota Board of Regents. These contents are solely the responsibility of the authors and do not necessarily represent the official views of the funding organizations. The study was conducted according to the guidelines of the University of Minnesota Institutional Animal Care and Use Committee (#2007-38247A approved on 08 August 2020). All data are contained within this manuscript and are available from the corresponding author upon reasonable request.

## REFERENCES

1. Chevalley T, Rizzoli R. Acquisition of peak bone mass. Best Pract Res Clin Endocrinol Metab. 2022;36(2):101616.

2. Demontiero O, Vidal C, Duque G. Aging and bone loss: new insights for the clinician. Ther Adv Musculoskelet Dis. 2012;4(2):61–76.

3. Sfeir JG, Drake MT, Khosla S, Farr JN. Skeletal Aging. Mayo Clin Proc. 2022;97(6):1194–208.

4. Wright NC, Looker AC, Saag KG, Curtis JR, Delzell ES, Randall S, et al. The recent prevalence of osteoporosis and low bone mass in the United States based on bone mineral density at the femoral neck or lumbar spine. J Bone Miner Res. 2014;29(11):2520–6.

5. Kannus P, Parkkari J, Niemi S, Palvanen M. Epidemiology of osteoporotic ankle fractures in elderly persons in Finland. Ann Intern Med. 1996;125(12):975–8.

6. Leibson CL, Tosteson AN, Gabriel SE, Ransom JE, Melton LJ. Mortality, disability, and nursing home use for persons with and without hip fracture: a population-based study. J Am Geriatr Soc. 2002;50(10):1644–50.

7. Magaziner J, Lydick E, Hawkes W, Fox KM, Zimmerman SI, Epstein RS, et al. Excess mortality attributable to hip fracture in white women aged 70 years and older. Am J Public Health. 1997;87(10):1630–6.

8. Magaziner J, Simonsick EM, Kashner TM, Hebel JR, Kenzora JE. Predictors of functional recovery one year following hospital discharge for hip fracture: a prospective study. J Gerontol. 1990;45(3):M101–7.

9. Riggs BL, Melton LJ, 3rd. The worldwide problem of osteoporosis: insights afforded by epidemiology. Bone. 1995;17(5 Suppl):505S–11S.

10. Clarke B. Normal bone anatomy and physiology. Clin J Am Soc Nephrol. 2008;3 Suppl 3(Suppl 3):S131–9.

11. Ramchand SK, Seeman E. The Influence of Cortical Porosity on the Strength of Bone During Growth and Advancing Age. Curr Osteoporos Rep. 2018;16(5):561–72.

12. Cosman F, Crittenden DB, Ferrari S, Khan A, Lane NE, Lippuner K, et al. FRAME Study: The Foundation Effect of Building Bone With 1 Year of Romosozumab Leads to Continued Lower Fracture Risk After Transition to Denosumab. J Bone Miner Res. 2018;33(7):1219–26.

13. Graeff C, Campbell GM, Pena J, Borggrefe J, Padhi D, Kaufman A, et al. Administration of romosozumab improves vertebral trabecular and cortical bone as assessed with quantitative computed tomography and finite element analysis. Bone. 2015;81:364–9.

14. Weivoda MM, Bradley EW. Macrophages and Bone Remodeling. J Bone Miner Res. 2023;38(3):359–69.

15. Sims NA, Martin TJ. Osteoclasts Provide Coupling Signals to Osteoblast Lineage Cells Through Multiple Mechanisms. Annu Rev Physiol. 2020;82:507–29.

16. Andreasen CM, El-Masri BM, MacDonald B, Laursen KS, Nielsen MH, Thomsen JS, et al. Local coordination between intracortical bone remodeling and vascular development in human juvenile bone. Bone. 2023;173:116787.

17. Liu B, Salgado OC, Singh S, Hippen KL, Maynard JC, Burlingame AL, et al. The lineage stability and suppressive program of regulatory T cells require protein O-GlcNAcylation. Nat Commun. 2019;10(1):354.

18. Wang H, Breed ER, Lee YJ, Qian LJ, Jameson SC, Hogquist KA. Myeloid cells activate iNKT cells to produce IL-4 in the thymic medulla. Proc Natl Acad Sci U S A. 2019;116(44):22262–8.

19. Wang H, Hogquist KA. CCR7 defines a precursor for murine iNKT cells in thymus and periphery. Elife. 2018;7.

20. Faltusova K, Chen CL, Heizer T, Bajecny M, Szikszai K, Paral P, et al. Altered Erythro-Myeloid Progenitor Cells Are Highly Expanded in Intensively Regenerating Hematopoiesis. Front Cell Dev Biol. 2020;8:98.

21. Mohamad SF, Gunawan A, Blosser R, Childress P, Aguilar-Perez A, Ghosh J, et al. Neonatal Osteomacs and Bone Marrow Macrophages Differ in Phenotypic Marker Expression and Function. J Bone Miner Res. 2021;36(8):1580–93.

22. McDonald MM, Khoo WH, Ng PY, Xiao Y, Zamerli J, Thatcher P, et al. Osteoclasts recycle via osteomorphs during RANKL-stimulated bone resorption. Cell. 2021;184(5):1330–47 e13.

23. Isojima T, Sims NA. Cortical bone development, maintenance and porosity: genetic alterations in humans and mice influencing chondrocytes, osteoclasts, osteoblasts and osteocytes. Cell Mol Life Sci. 2021;78(15):5755–73.

24. Molstad DHH, Zars E, Norton A, Mansky KC, Westendorf JJ, Bradley EW. Hdac3 deletion in myeloid progenitor cells enhances bone healing in females and limits osteoclast fusion via Pmepa1. Sci Rep. 2020;10(1):21804.

25. Helming L, Winter J, Gordon S. The scavenger receptor CD36 plays a role in cytokine-induced macrophage fusion. J Cell Sci. 2009;122(Pt 4):453–9.

26. Ambrosi TH, Marecic O, McArdle A, Sinha R, Gulati GS, Tong X, et al. Aged skeletal stem cells generate an inflammatory degenerative niche. Nature. 2021;597(7875):256-62.

27. Ucer S, Iyer S, Kim HN, Han L, Rutlen C, Allison K, et al. The Effects of Aging and Sex Steroid Deficiency on the Murine Skeleton Are Independent and Mechanistically Distinct. J Bone Miner Res. 2017;32(3):560–74.

28. Yahara Y, Ma X, Gracia L, Alman BA. Monocyte/Macrophage Lineage Cells From Fetal Erythromyeloid Progenitors Orchestrate Bone Remodeling and Repair. Front Cell Dev Biol. 2021;9:622035.

29. Askmyr MK, Fasth A, Richter J. Towards a better understanding and new therapeutics of osteopetrosis. Br J Haematol. 2008;140(6):597–609.

30. Jacome-Galarza CE, Percin GI, Muller JT, Mass E, Lazarov T, Eitler J, et al. Developmental origin, functional maintenance and genetic rescue of osteoclasts. Nature. 2019;568(7753):541-5.

31. Novak S, Roeder E, Kalinowski J, Jastrzebski S, Aguila HL, Lee SK, et al. Osteoclasts Derive Predominantly from Bone Marrow-Resident CX3CR1(+) Precursor Cells in Homeostasis, whereas Circulating CX3CR1(+) Cells Contribute to Osteoclast Development during Fracture Repair. J Immunol. 2020;204(4):868–78.

32. Xia Y, Inoue K, Du Y, Baker SJ, Premkumar Reddy E, Greenblatt MB, et al. TGFbeta reprograms TNF stimulation of macrophages towards a non-canonical pathway driving inflammatory osteoclastogenesis. Nat Commun. 2022;13(1):3920.

33. Yahara Y, Barrientos T, Tang YJ, Puviindran V, Nadesan P, Zhang H, et al. Erythromyeloid progenitors give rise to a population of osteoclasts that contribute to bone homeostasis and repair. Nat Cell Biol. 2020;22(1):49–59.

34. Kevorkova O, Martineau C, Martin-Falstrault L, Sanchez-Dardon J, Brissette L, Moreau R. Low-bone-mass phenotype of deficient mice for the cluster of differentiation 36 (CD36). PLoS One. 2013;8(10):e77701.

35. Koduru SV, Sun BH, Walker JM, Zhu M, Simpson C, Dhodapkar M, et al. The contribution of cross-talk between the cell-surface proteins CD36 and CD47-TSP-1 in osteoclast formation and function. J Biol Chem. 2018;293(39):15055–69.

36. Weivoda MM, Ruan M, Hachfeld CM, Pederson L, Howe A, Davey RA, et al. Wnt Signaling Inhibits Osteoclast Differentiation by Activating Canonical and Noncanonical cAMP/PKA Pathways. J Bone Miner Res. 2016;31(1):65–75.

37. Glass DA, 2nd, Bialek P, Ahn JD, Starbuck M, Patel MS, Clevers H, et al. Canonical Wnt signaling in differentiated osteoblasts controls osteoclast differentiation. Dev Cell. 2005;8(5):751–64.

38. Andersen TL, Sondergaard TE, Skorzynska KE, Dagnaes-Hansen F, Plesner TL, Hauge EM, et al. A physical mechanism for coupling bone resorption and formation in adult human bone. Am J Pathol. 2009;174(1):239–47.

39. Eriksen EF, Gundersen HJ, Melsen F, Mosekilde L. Reconstruction of the formative site in iliac trabecular bone in 20 normal individuals employing a kinetic model for matrix and mineral apposition. Metab Bone Dis Relat Res. 1984;5(5):243–52.

40. Lassen NE, Andersen TL, Ploen GG, Soe K, Hauge EM, Harving S, et al. Coupling of Bone Resorption and Formation in Real Time: New Knowledge Gained From Human Haversian BMUs. J Bone Miner Res. 2017;32(7):1395–405.

41. Bouxsein ML, Boyd SK, Christiansen BA, Guldberg RE, Jepsen KJ, Muller R. Guidelines for assessment of bone microstructure in rodents using micro-computed tomography. J Bone Miner Res. 2010;25(7):1468–86.

42. Mattson AM, Begun DL, Molstad DHH, Meyer MA, Oursler MJ, Westendorf JJ, et al. Deficiency in the phosphatase PHLPP1 suppresses osteoclast-mediated bone resorption and enhances bone formation in mice. J Biol Chem. 2019;294(31):11772–84.

43. McGee-Lawrence ME, Razidlo DF. Induction of fully stabilized cortical bone defects to study intramembranous bone regeneration. Methods Mol Biol. 2015;1226:183–92.

